# MjCRSP, a small cysteine-rich effector protein from *Meloidogyne javanica* suppresses plant immunity

**DOI:** 10.1101/2025.09.18.677000

**Authors:** Teresia Macharia, Lucy Novungayo Moleleki

**Author notes:** Corresponding author: Lucy Moleleki.

## Abstract

*Meloidogyne javanica* is one of the most important root-knot nematode (RKN) species causing considerable economic losses globally. Effectors which are secreted nematode molecules play an important role in mediating nematode interaction with their hosts. Specifically, small cysteine rich (SCR)- secreted proteins found across many taxa, such as bacteria oomycete and fungi are of interest for this particular study. Towards this end, we identified over 600 SCR in the genome of *M. javanica*, with a size range between 100 and 300 amino acids (aa) with 4 – 8 cysteine residues. We selected one of these proteins, which we named MjCRSP for further characterisation. MjCRSP was predicted to have a signal peptide at the N-terminus end and a putative nuclear localization signal (NLS) in the C-terminus region. Using *Agrobacterium*-mediated transient expression system in *Nicotiana benthamiana*, we show that this protein inhibits elicitin infestin1 (INF1)-triggered cell death and immune responses triggered by Flg22 peptide. Furthermore, subcellular localization showed that MjCRSP localizes in the cytoplasm. This study demonstrated that MjCRSP is *M. javanica* effector protein that modulates plant immune responses during RKN-host interactions.

## Introduction

Root-knot nematodes belong to a significant group of sedentary endoparasites, the genus, *Meloidogyne*. Like other phytopathogens, RKNs release an arsenal of proteinaceous molecules, known as effectors, at the host-nematode interface, which are essential for host colonization (Jagdale et al., 2021). The pattern-triggered immunity (PTI) and effector-triggered immunity (ETI) of plants, which serve as an adaptation against parasites, work in concert to fight pathogen invasion (Bentham et al., 2020, Lu & Tsuda, 2021). It is also worth noting that PTI, which is activated by nematode-associated molecular patterns or danger signals detected by cell-surface receptors and it is involved in plant defence against RKN. Consequently, plant-parasitic nematodes (PPNs) have effectively evolved effectors to overcome both PTI and ETI (Goode & Mitchum, 2022, Siddique et al., 2022).

In a host-biotrophic pathogen system, the host is impacted by pathogen-effector molecules. When the corresponding host resistance proteins recognize the pathogen effector molecules, this leads to ETI, which is accompanied by programmed cell death (PCD) and the activation of a plethora of plant defense responses. ETI response culminates in a resistant or incompatible interaction, which is intended to avert further colonization of a biotrophic pathogen (Bentham et al., 2020), hence biotrophs, such as RKNs, avoid or suppress plant defense (Haegeman *et al*., 2012). This notion is well supported by several research studies utilizing *Agrobacterium-*mediated transient system showing that RKN effectors largely do not function as elicitors of PCD, but rather as suppressors of induced cell death by various known elicitors (Niu et al., 2016, Naalden et al., 2018, Kamaruzzaman et al., 2022, Zhuo et al., 2017, Chen et al., 2017, Song et al., 2021b, Hu et al., 2022). Although the mechanism of RKN effectors as cell death suppressors is unknown, we can speculate that the interaction between the RKN effector and its host cognate protein may help the nematode avoid detection or inhibit plant defensive response activation, which could cause a resistant response.

Plant-parasitic nematodes have a significant impact on the global food supply. Nematicides were previously the frontier for managing nematode diseases, but their adverse human and environmental implications have required agricultural specialists to explore alternatives that contribute to food security (Coyne et al., 2018). A crucial aspect of pest management is to understand how these parasites penetrate, target, and manipulate the host (Zhang et al., 2020). Therefore, understanding basic effector biology is vital for tackling the threat emanating from these parasitic infestations (Varden *et al*., 2017). In keeping with this, the subject of effector biology has become increasingly important in recent years, with an emphasis on understanding how pathogen effectors promote successful infection. In addition to advancing our knowledge of plant pathogenesis, the information gleaned from effector research biology may also open new possibilities for developing measures to control pests and pathogens.

Research studies focusing on the discovery and functional characterization of RKN-effector biology is expanding enormously. The development of low-cost sequencing and molecular biology technology has greatly contributed to research focused on identifying functional effectors encoded by pioneer genes in nematodes (Vieira & Gleason, 2019). Nematode effectors engage in various functions such as immunomodulators that shield the parasite from plant defenses such as PCD, cell wall degrading and modifying enzymes that contribute to nematode invasion and migration, plant reprogramming modulators that facilitate in feeding site formation and maintenance, and host metabolism modulators that enable nematode nutrient acquisition (Niu et al., 2016, Haegeman et al., 2012).

Small cysteine-rich (SCR)-secreted proteins have garnered significant attention among plant pathologists (Stergiopoulos & de Wit, 2009). This is because, besides being found across many taxa such as bacteria, oomycetes, fungi, and plants; characterization of SCR has revealed several elicitors, virulence, and avirulence factors or effectors, such as VmE02 from *Valsa mali* (Nie *et al*., 2019), PnSCR82 from *Phytophthora nicotianae* (Wang *et al*., 2022), RsMf8HN from *Rhizoctania solani* (Niu *et al*., 2023), and PmSCR1 from *Pythium myriotylum* (Wang *et al*., 2023). Furthermore, Cys residues in SCR-secreted proteins generate internal disulfide connections, which improve protein stability in the presence of proteases present in plant apoplasts. The possibility that SCR proteins are a class of secreted proteins that interact with host proteins (Niu et al., 2023) is a plausible rationale for the surge in SCR protein research.

Comparable to other pathogen genomes, one to several hundred SCR-secreted proteins are also encoded by PPN genomes (Petitot et al., 2016). BxSCD1 from *Bursaphelenchus xylophilus* (Wen *et al*., 2021), MiSGCR1 (Nguyen et al., 2018), and MiCTL1 (Zhao et al., 2021) both from *M. incognita* are good examples of functionally characterized SCR effectors from PPNs. Small cysteine-rich proteins that are shown to be secreted have been proposed as plausible candidate effector proteins that play an important role during host-pathogen interactions (Wang *et al*., 2020). However, SCR proteins’ mechanism and function in plant-nematode interactions require further exploration.

In this study, using comparative analysis, we identified a small cysteine-rich secreted protein encoded in *M. javanica* genome denoted as *Meloidogyne javanica* cysteine-rich secreted protein1 (hereafter, MjCRSP). Here we present evidence that MjCRSP is a bona fide effector, is localized in the nematode amphidial secretory glands, and suppresses plant defense responses.

### Experimental procedures

#### Nematode culture and plant growth conditions

A pure population of *M. javanica* species was cultured on tomato plants (*Solanum lycopersicum* cv Floradade) at 25 °C in a glasshouse. To hatch pre-parasitic J2s (pre-J2s), egg masses on root galls were handpicked and hatched over sterile water at 28 °C in sterile petri dishes for 48-72 h. The pre-J2s were collected for use in the root infection assays or for extraction of nucleic acids. Total genomic DNA from *M. javanica*-infested roots was isolated using MasterPure (DNA extraction kit) using the manufacturer’s instructions.

#### *Meloidogyne javanica* effector identification through data mining and *in-silico* analysis

Comparative BLAST analyses were conducted between a database of *M. incognita* parasite-specific effectors by Grynberg et al. (2020) and a database of *M. javanica* secreted proteins by Macharia et al. (2023) to identify a putative *M. javanica* effector. *Meloidogyne javanica*_Scaff7153g046964 effector coding gene, a homolog of Minc3s05519g38359 in *M. incognita,* highly expressed during endophytic stages (Da Rocha et al., 2021), was selected for further functional characterization. Homologous sequences of MjCRSP in other RKN species were searched by querying the NCBI, WormBase ParaSite, and *Meloidogyne* genomic resources using the protein BLAST tool. Multiple sequences were aligned using Muscle software (Edgar, 2004); and the phylogenetic tree was constructed with NGPhylogeny.fr (Lemoine *et al*., 2019).

In addition, we used Pairwise identity matrices generated using Sequence Demarcation Tool Version 1.2 software (Muhire *et al*., 2014). HHpred (PDB_mmCIF70_17_Apr as the structural/domain database) (https://toolkit.tuebingen.mpg.de/tools/hhpred) (Zimmermann *et al*., 2018) was used for predicting remote homology of known structure.

#### Plasmid construction and transient expression analysis

The candidate MjCRSP effector was cloned without its predicted signal peptide. For cloning, the sequence was amplified from *M. javanica* genomic DNA using PCR with gene-specific primers flanking Gateway^®^ recombination sites (**Table S1**). The PCR products were purified and then recombined into pDONR201 using BP clonase reaction to generate entry clones. The entry clones were shuttled into destination vectors pB7WGF2-GFP using LR recombination reaction. The constructs were confirmed using colony PCR and DNA sequencing and then transformed into *Agrobacterium tumefaciens* GV3101 strain through electroporation.

The transformed *A. tumefaciens* strains were cultured overnight with shaking in yeast-extract and beef (YEB) medium at 28 °C with selective antibiotics. The bacteria cultures containing the constructs were pelleted by centrifugation at 5000g for 15 mins and resuspended in infiltration buffer (10mM MES, 10mM MgCl_2,_ and 200mM Acetosyringone, pH 5.5). For Agroinfiltration, suspended *Agrobacterium* cells were adjusted to a final optical density at 600 nm of 0.5, and the cell suspension was infiltrated into plant leaves using a 1-ml syringe without a needle. Before infiltration, the bacterial suspensions were incubated at room temperature in the dark for at least 2-3 hours.

To determine the cell death-inducing or -inhibiting activity of the MjCRSP candidate effector protein, the P35s-MjCRSP^-SP^ and p35s eGFP control vector were agroinfiltrated into four-week *N. benthamiana* leaves 24 h before INF1/ Avr3a: R3a constructs were infiltrated at the same site. INF1, GFP, Avr3a, and R3a were used as controls. Transient co-expression of R3a and Avr3a were prepared as described by (Bos *et al*., 2006). Cell death symptoms were monitored visually up to 7 days post infiltration. Cell death quantification was performed using LeafQuant-T3S, a Matlab-based image-processing package R2021a (Ramachandran *et al*., 2017). The experiments were done in triplicates with five plants per assay.

#### Subcellular localization analysis

The subcellular localization of MjCRSP was determined by transient expression in *N. benthamiana* leaves. The constructs carrying P35s-MjCRSP^-SP^, and P35s-eGFP were agroinfiltrated into *N. benthamiana* leaves and harvested at 2 dpi for imaging. Images of the fluorescent protein constructs in *N. benthamiana* epidermal cells were obtained using a Zeiss LSM 510 confocal microscope (Carl Zeiss). The excitation wavelength was 488nm for eGFP and DAPI, respectively.

#### Reactive oxygen species and callose deposition assays

Detection of H_2_O_2_ in leaves was carried out using the 3,3-diaminobenzidine (DAB) staining method described previously (Daudi & O’Brien, 2012). Agrobacteria carrying P35s-MjCRSP^- SP^ and p35s-eGFP constructs were transiently expressed in *N. benthamiana* followed by Flg22 peptide inoculation at 48 hours post infiltration (hpi) incubated for 1 hr. Leaf segments were incubated in DAB staining solution overnight to allow DAB uptake and reaction with H_2_O_2_. Solutions were kept under dark conditions. After incubation, leaves were decolorized in boiling (∼80 °C) destaining solution (ethanol: acetic acid: glycerol 3:1:1) for 20 min and transferred into a solution containing water and 20% glycerol. Leaf segments were placed in filter paper to remove the excess glycerol solution, scanned (Epson L3150) and images were analyzed using ImageJ software.

Callose deposition was visualized using the aniline blue staining approach as described previously (Schenk & Schikora, 2015). *Agrobacterium* constructs with P35s-MjCRSP^-SP^ and p35s eGFP were transiently expressed in *N. benthamiana* followed by Flg22 peptide inoculation at 48 hpi. Leaf disks expressing the various constructs were soaked overnight in a destaining solution (ethanol: acetic acid 3: 1) at room temperature with gentle shaking. Thereafter, the destained leaves were incubated in 0.01% aniline blue in 150mM K_2_HPO_4_ at room temperature subsequently imaged using a fluorescence microscope and analyzed with ImageJ software. Both ROS and callose deposition assays were performed at least three independent times with leaves obtained from 3 to 5 plants.

#### Fluorescence in situ hybridization (FISH) of MjCRSP

Electropermeabilization-based fluorescence in situ hybridization of whole-mount *M. javanica* specimens was conducted based on (Ruark-Seward *et al*., 2019) protocol. The DNA (22-mer) probe complementary to the target sequence of MjCRSP was designed using Probe melt software and checked for specificity using blast analysis software. The probes were custom-synthesized by Whitehead Scientific (Pty) Ltd. The probe was covalently linked at its 5’ end; ATTO 647N 5’- CAAGTATCATCCTCTCCACTAC-3’. Roughly 1, 000 freshly hatched pre-parasitic J2s *M. javanica* stages were collected from infected roots by root blending and sieving. The samples were purified from root debris by filtering through sieves with 38-µm pores. The J2s were washed with M9 buffer to remove contamination followed by rinsing in sterile water. The samples were resuspended in 50 μl sterile water, and the nematode suspension was pipetted into an electroporation cuvette and the 150μm of the probe [(1.5 μl of 100 μm concentrated probe in TE (10 mM Tris-HCl, 1 mM EDTA, pH 7.4)] was added.

Nematodes suspended in probe solution were electroporated on a BIO-RAD Electroporation apparatus with a single-pulse square wave at 200 V for 20 ms. After electroporation, the nematodes were hybridized overnight at 46°C in 100 μl of prewarmed hybridization buffer in the dark on a heating block. Following hybridization, three incubations were performed at 50°C in 100 μl of pre-warmed hybridization wash buffer to remove the excess probe. Final centrifugation was conducted at 8,000×g for 2 min, and excess liquid was removed from the nematode pellet. The stained samples were submerged in 20μl of hybridization wash buffer and were transferred to a slide sealed with nail polish and viewed under Zeiss LSM 510 confocal microscope (Carl Zeiss). Detection specificity was confirmed using *M. javanica* exposed to sterile water as a control.

#### Statistical analysis

Mini tab was used to analyze the experimental data and the Tukey method at 95% (*P* ≤ 0.05) confidence was used to determine statistical differences.

## Results

### Identification of secreted small cysteine-rich (SCR) proteins encoded in the *M. javanica* genome

Small cysteine-rich proteins found across all life forms are important for, amongst other roles, modulating plant defences. A large number of SCR have been predicted in the genome of some *Meloidogyne species* (Petitot et al., 2016). Here, our goal was to identify SCRs encoded in *M. javanica* genome, ascertain the abundance of cysteine residues and establish the conserved domains represented in *M. javanica* SCR proteins. Towards this end, we evaluated the presence and abundance of cysteine residues in *M. javanica* predicted secretome reported recently by Macharia et al., 2023 wherein 2,848 of the 5,258 predicted proteins designated as classically secreted proteins had four cysteine residues in the mature peptide. Most MjSCRs are between 100 and 300 amino acids with 4-8 cysteine residues (**Figure 1a and b**). This finding implies that a high cysteine abundance is present in small secreted proteins, similar to what has been described for fungal effectors (Sperschneider *et al*., 2015).

**Figure 1:**
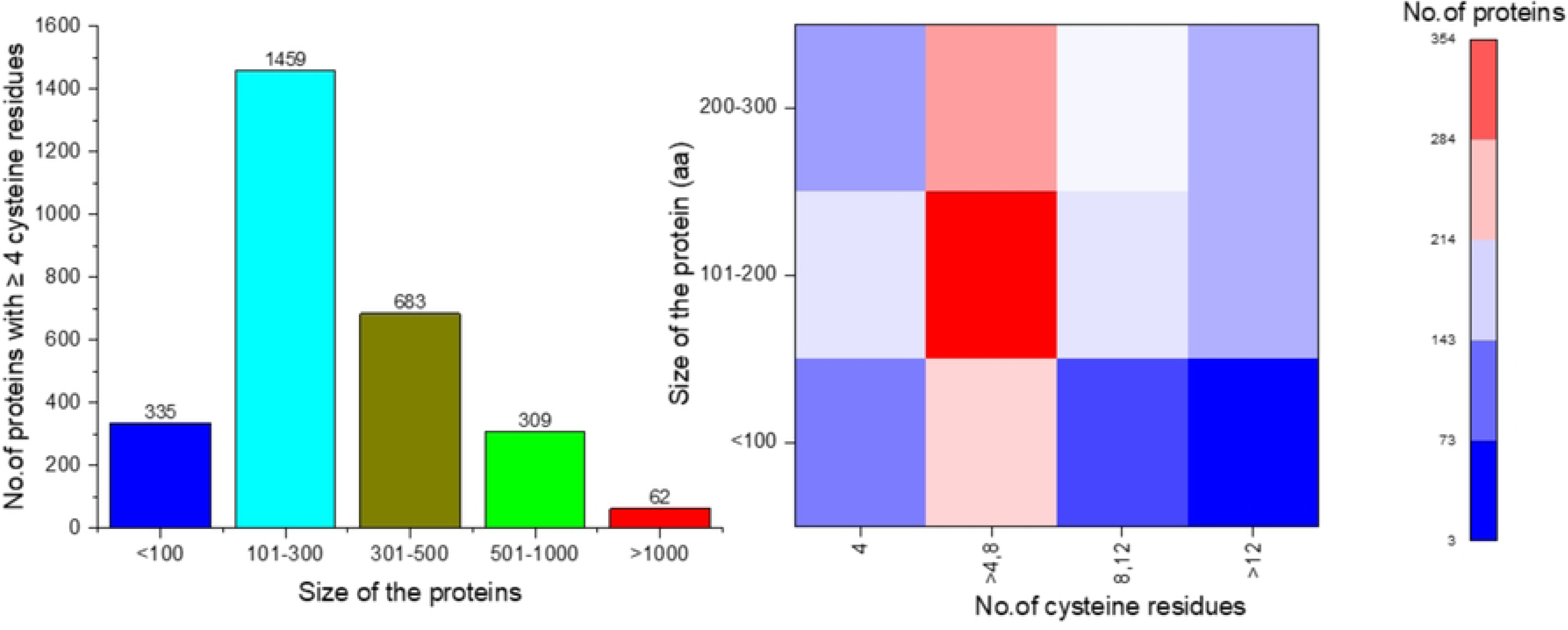
A summary of small, secreted cysteine-rich protein s encoded in *M. javanica* genome. (a) Analysis of the number of secreted *M. javanica* proteins with 4 or more cysteine residues in the mature peptide. (b) Distribution by protein size and cysteine content.

Using the NCBI conserved domain database, 632 proteins of the small-cysteine-secreted proteins (1794 proteins with sequence lengths between 100-300 aa) were mapped into 189 families. The top represented families include some known functionally characterized effectors from RKN such as those within the pectate lyase family (Chen et al., 2021), C-type lectin domain (Zhuo et al., 2019, Zhao et al., 2021), and transthyretin-like protein (Lin et al., 2016).

### MjCRSP is a small-secreted protein with a cysteine-rich region and is conserved within root-knot nematodes

The open reading frame of MjCRSP (Mjav1s07153g046964, European Nucleotide archive accession CEWN01000000.1) contains 258 nucleotides which encode for 85 amino acid (aa) peptide. Bioinformatic analysis revealed the existence of a 22 amino acid signal peptide at the N-terminus region, where the cleavage site was predicted between amino acid positions A^22^ and A^23^(**Figure S1**) Additionally, LOCALIZER (Sperschneider et al., 2017) predicted the presence of a nuclear localisation signal (NLS) in MjCRSP (KVKK), likely mediating protein transport from the cytoplasm into the nucleus (Lu et al., 2021). The protein has a molecular weight of 9227.73 and an isoelectric point of 8.63. Moreover, no transmembrane topology was detected by Phobius (Käll et al., 2004) and TMHMM2.0 (Krogh, 2006). Using Deeploc and LocTree3 (Goldberg *et al*., 2012) MjCRSP was predicted as an extracellular protein (Gene ontology term extracellular region, GO term ID: <u>GO:0005576</u>) suggesting that the host-parasite interface might be the site of this protein action.

Out of the 85 aa found in MjCRSP, 12 aa are cysteine residues which account for 14.11% of the full protein sequence (**Figure 2a**). Cysteine-rich proteins typically feature structural homologies based on Cys residues, such as the formation of disulfide bonds, which play an important role in folding and, consequently, the chemical stability of the molecule (Petitot et al., 2016). MjCRSP protein function prediction using the HHpred server Zimmermann et al., 2018 showed a top significant hit (1.7e-8 and 98.64 % probability) identified as A7RZ3_A Xt3a, a double inhibitor cysteine knot toxin (ICK) (Maxwell *et al*., 2023). A7RZ3_A Xt3a encodes a bivalent toxin from remipede, *Xibalbanus tulumensis* that target ryanodine receptors (Maxwell *et al*., 2023) which are huge ion channels that responsible for ca^2+^ releases from sacro/endoplasmic reticulum and thus control many Ca^2+^-dependent processes within the cell (Van Petegem, 2012). Ion channels are the fastest cellular signaling elements, thus it is not surprising that many venomous species have evolved to target these proteins to induce a rapid physiological response (Maxwell et al., 2023). Various biological roles have been linked to these peptides including enzymatic inhibition, predation, and defense (Zhao et al., 2018). An inhibitor cystine knot is an ultra-stable structural motif containing three disulfide bridges. They are extensively dispersed in arthropods, plants, fungi, animals, and certain viruses (Craik *et al*., 2001, Zhao *et al*., 2018, Park *et al*., 2014). The structural domain is located at A^21^- A^81^ where the cysteine-rich region profile lies (**Figure 2a**).

**Figure 2:**
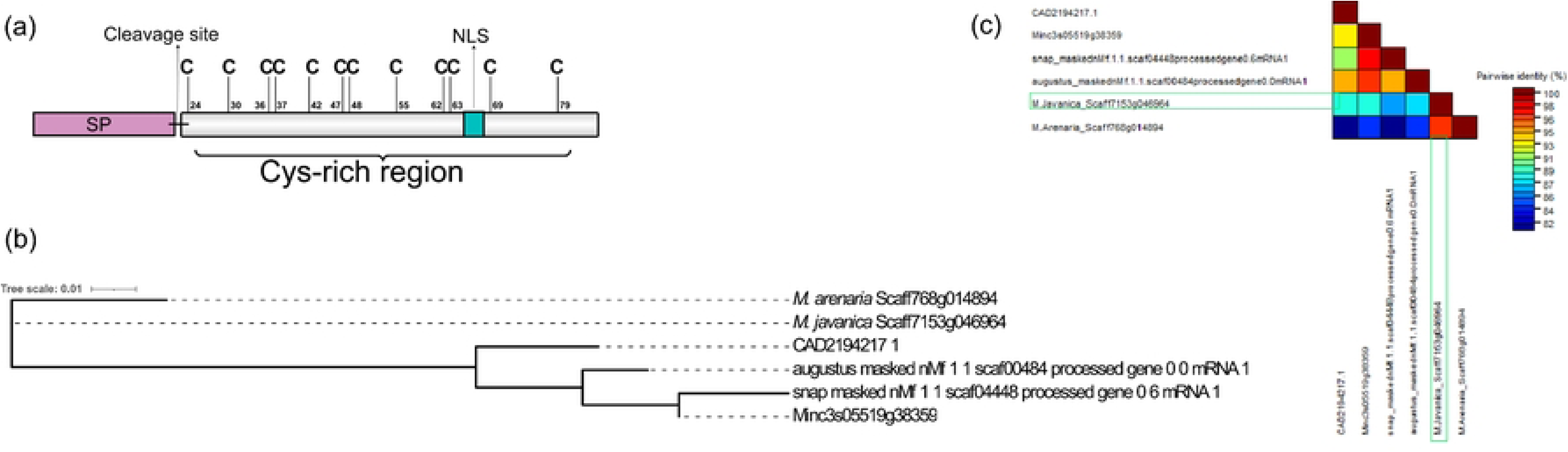
Sequence analysis of MjCRSP and its homologs from other root-knot nematode species. a) A schematic representation of the primary sequence of the MjCRSP. (b) Pairwise sequence identity matrices of amino acid sequences were generated using Sequence Demarcation Tool version 1.2 software. (c) The Phylogeny of MjCRSP and its homologous sequences from different RKN species were generated by the Phylogeny.fr web service. The RKN protein of interest in this study is marked with an asterisk. Abbreviation: SP-signal peptide, NLS-nuclear localization signal.

After searching several protein databases, MjCRSP homologs were only detected in RKN species. Pairwise comparison of these protein sequences showed that MjCRSP and its homologs have a high sequence similarity to each other ranging from 82-100% (**Figure 2b and 2c**). All the homologs, except for the *M. arenaria* homolog, were predicted to have a signal peptide at the N-terminus with a cleavage site between A^22^ and A^23^. This suggests that MjCRSP is conserved in *Meloidogyne* species and might be adapted to play an indispensable role in their pathogenicity process.

### MjCRSP suppresses plant immunity

Here, we employed the *Agrobacterium* transient transformation assay to evaluate the putative function of this RKN effector in *N. benthamiana*, we tested whether MjCRSP could induce or inhibit PCD triggered by the well-known INF1, a PAMP from *Phytophthora infenstans* or the AVRa/R3a cognate ETI induced cell death. In these assays, INF1 and AVRa/R3a were used as positive controls and green fluorescent protein (GFP hereafter, empty vector- (EV) co-expressed with INF1 and/or AVRa/R3a as negative control. As shown in **Figure 3 a1 and b1** MjCRSP did not induce cell death at 4 days after infiltration suggesting that MjCRSP evades plant immune responses. The expression of INF1 or AVRa/R3a with MjCRSP showed that MjCRSP blocks the INF1-mediatedcell death when compared to empty vector (**Figure 3 a2 and b2**), but it does not inhibit the ETI-induced cell death (**Figure 3 a3 and b3**).

**Figure 3:**
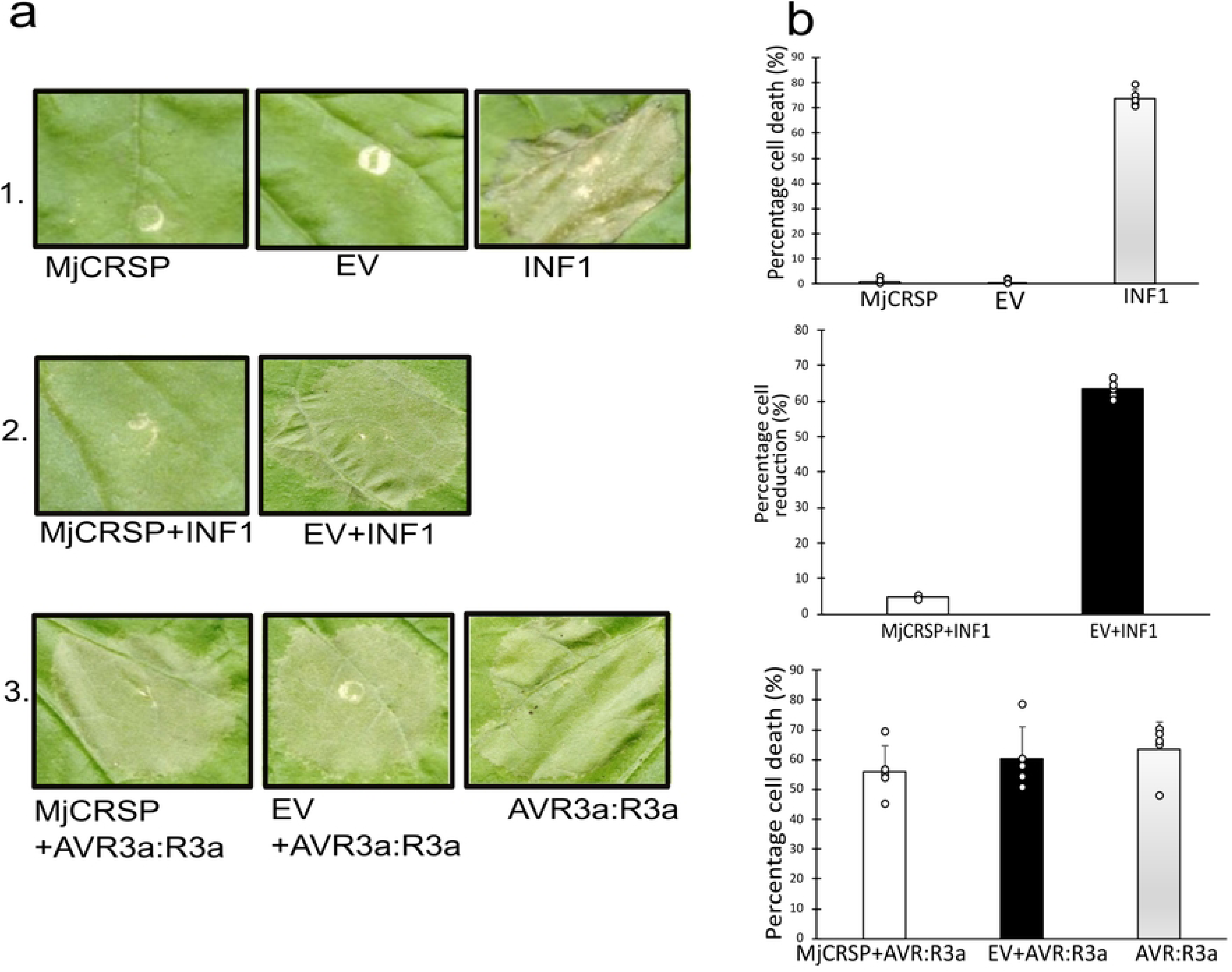
MjCRSP inhibits immunity-associated cell death in *Nicotiana benthamiana*. (a1, a2 and a3) Representative cell death symptoms on *N. benthamiana* leaves infiltrated with MjCRSP, empty vector, INF1, and AVR3a: R3a co-infiltrations. (b1, b2, b3) Quantification of cell death symptoms using LeafQuant-T3S, a Matlab-based image-processing package R2021a. MjCRSP did not cause cell death in *N. bethamiana* leaves. (a2 and b2) MjCRSP suppresses INF1-triggered cell death in *N. benthamina* leaves. (3a and 3b) MjCRSP did not inhibit cell death induced by AVR3a/R3a constructs. *Agrobacterium tumefaciens* construct carrying MjCRSP was infiltrated into the leaves of *N. benthamiana* 24 h before infiltration with cells expressing INF1 and/or AVR3a/R3a. The EV was used as a negative control. At least five separate biological repetitions were used for each experiment. The experimental data was analyzed using Minitab, and the differences were found using Tukey’s test at P≤ 0.05.

### Transient expression of MjCRSP inhibits immune responses triggered by the Flg22 PAMP in *N. benthamiana* leaves

Having established that MjCRSP inhibits INF1-mediated cell death, we hypothesized that MjCRSP may be an essential nematode virulence effector, likely interfering with PAMP-triggered immune responses such as those triggered by Flg22. Towards this end, we evaluated the buildup of ROS and callose deposition in *N. benthamiana* plants to confirm whether MjCRSP interferes with plant immunity through the responses mentioned above. To test this, we infiltrated *N. benthamiana* leaves with MjCRSP or an empty vector control. At 2 days post infiltration, we inoculated the leaves with 100 µm Flg22 which induces the immune responses. As shown in **Figures 4a and b**, ROS buildup and callose deposition in leaves expressing MjCRSP+Flg22 were significantly lower than in leaves expressing the EV+Flg22 control. These findings imply that MjCRSP expression in *N. benthamiana* suppresses plant immunological responses induced by the Flg22 PAMP.

**Figure 4:**
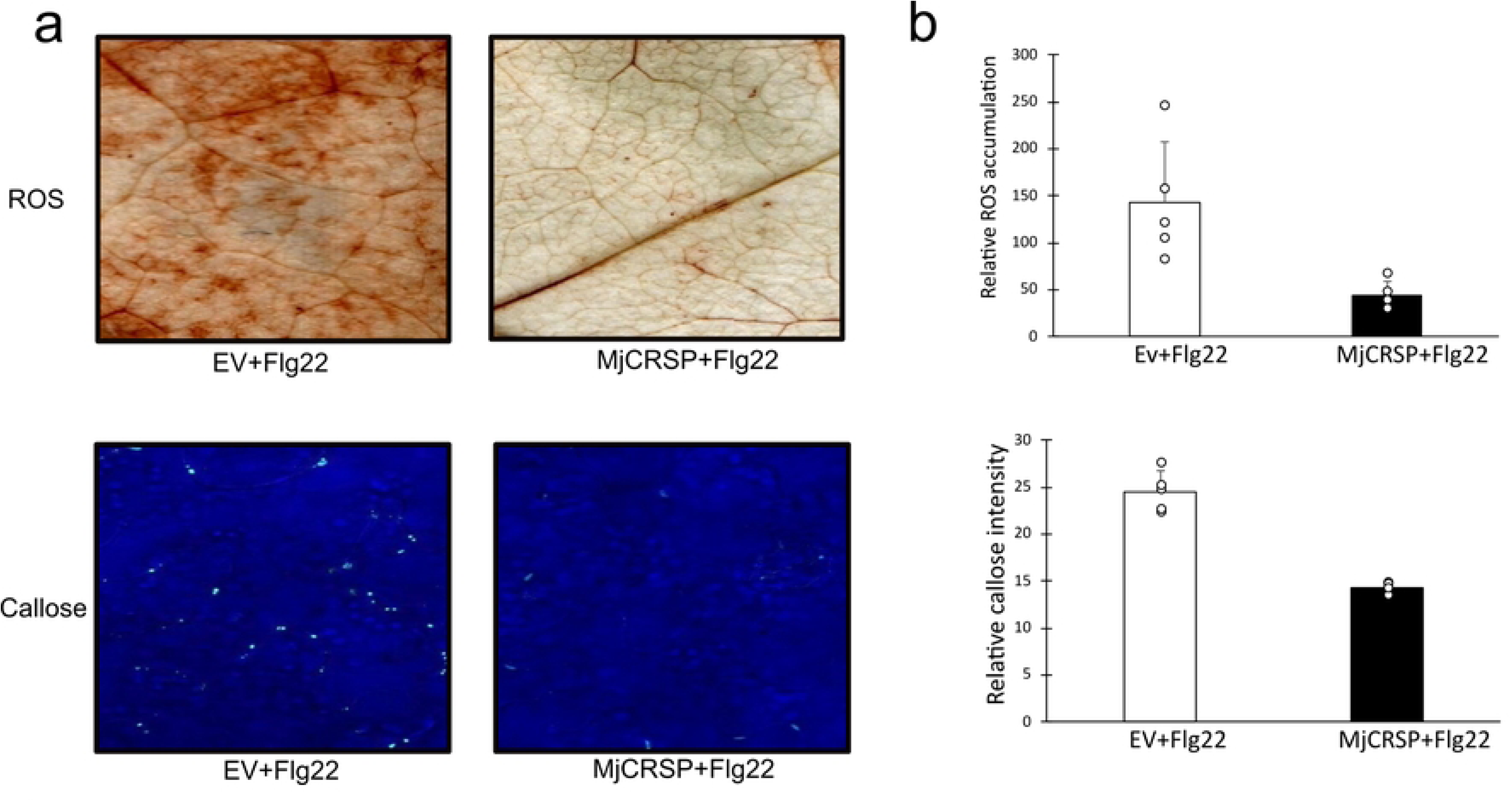
Co-expression of MjCRSP and Flg22 peptide in *Nicotiana benthamiana* suppresses PAMP-triggered immunity. (a) Accumulation of ROS, and deposition of callose. (b) Quantification of qualitative ROS, and callose intensity with ImageJ software. At least five separate biological repetitions were used for each experiment. The experimental data was analyzed using Minitab, and the differences were found using Tukey’s test at P≤ 0.05.

### Subcellular localization of MjCRSP *N. benthamiana* cells

Once they are secreted outside the parasite body, RKN effectors are delivered to various host cellular compartments including the apoplast (Tian et al., 2019), cytoplasm (Niu et al., 2016, Song et al., 2021a), or nucleus (Song et al., 2021b). Given that our earlier in silico prediction identified a potential NLS, proceeded to assess the subcellular localization of MjCRSP. Towards this end, we transiently expressed MjCRSP without its native SP in *N. bethamiana* leaves. At 2 days post infiltration, confocal microscopy analysis revealed that the fluorescence of the GFP-tagged MjCRSP protein was present in the cytoplasm and the nucleus (**Figure 5a**),

**Figure 5:**
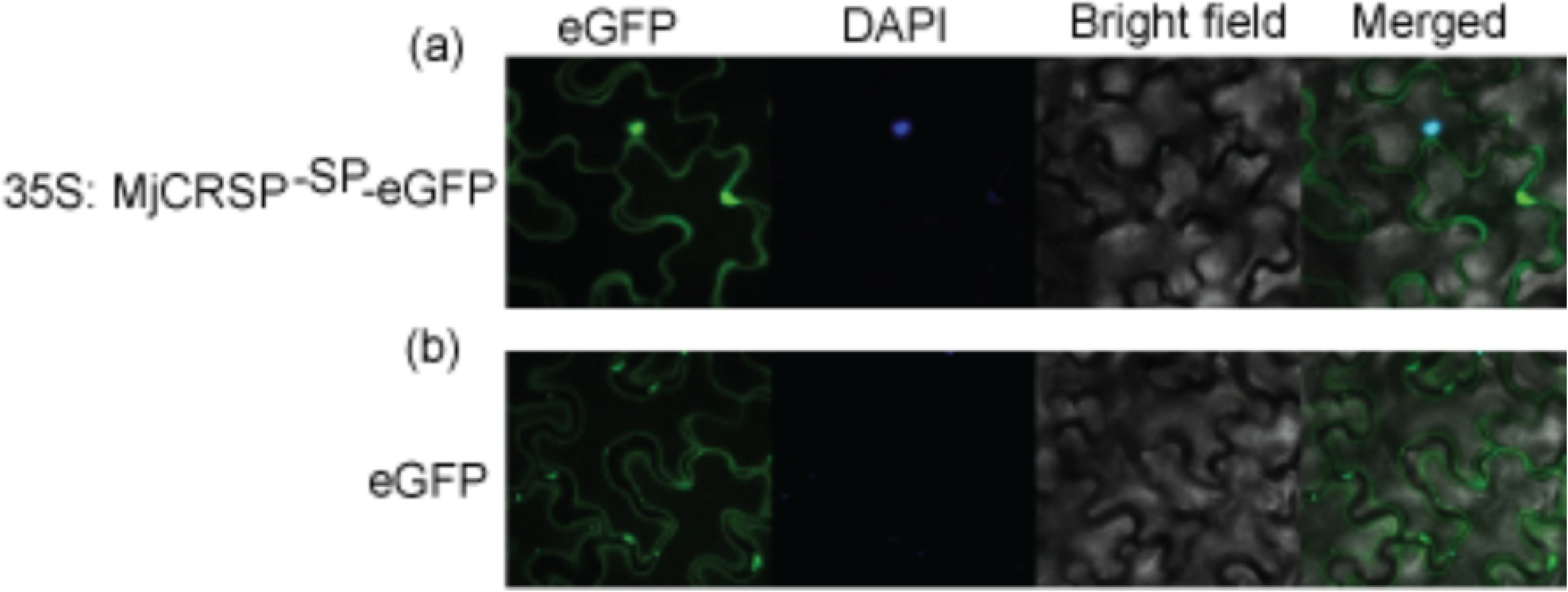
MjCRSP subcellular localization in *N. benthamiana* epidermal cells. Green fluorescent protein (eGFP)-tagged protein was transiently expressed in *N. benthamiana* leaves to assess the subcellular location of MjCRSP. Confocal microscopy images taken 2 days post-infiltration showed the subcellular localization of MjCRSP in the cytoplasm and nucleus (confirmed with bltable of *N. bethamiana* epidermal cells. An empty vector with eGFP was used as a control.

Nuclear counter staining (in blue) confirmed that the effector localizes to the nucleus. On the other hand, eGFP was largely found in the cytoplasm (**Figure 5b**). It is noteworthy that the MjCRSP: GFP fusion protein has a size of just 36 kDa, which is sufficiently small to permit passive diffusion into the nucleus. Whether the uptake of the effector into the nucleus is indeed passive or otherwise, remains to be fully established.

### MjCRSP is localized in the amphids: a major sensory gland

Fluorescence in situ hybridization was used to evaluate the spatial localization of MjCRSP in *M. javanica* tissue. Pre-parasitic J2s nematode samples were hybridized with MjCRSP gene-specific probe conjugated with the fluorescent dye ATTO 647N. A strong signal was observed in the secretory glands located in the anterior region of the nematode, the amphids (**Figure 6a**). No signal was visible on the negative control, nematodes hybridized without the probe (**Figure 6b**). Overall, the dorsal gland and sub ventral glands are the two largest best-studied secretory glands where plant-parasitic nematode effectors are mostly produced. However, secreted proteins can also originate from other organs, including the hypodermis, intestines, amphids, and phasmids (Vieira & Gleason, 2019; Nguyen et al., 2018; Wen et al., 2021; Jagdale et al., 2021). Thus, while this result was somewhat unexpected, it is not improbable.

**Figure 6:**
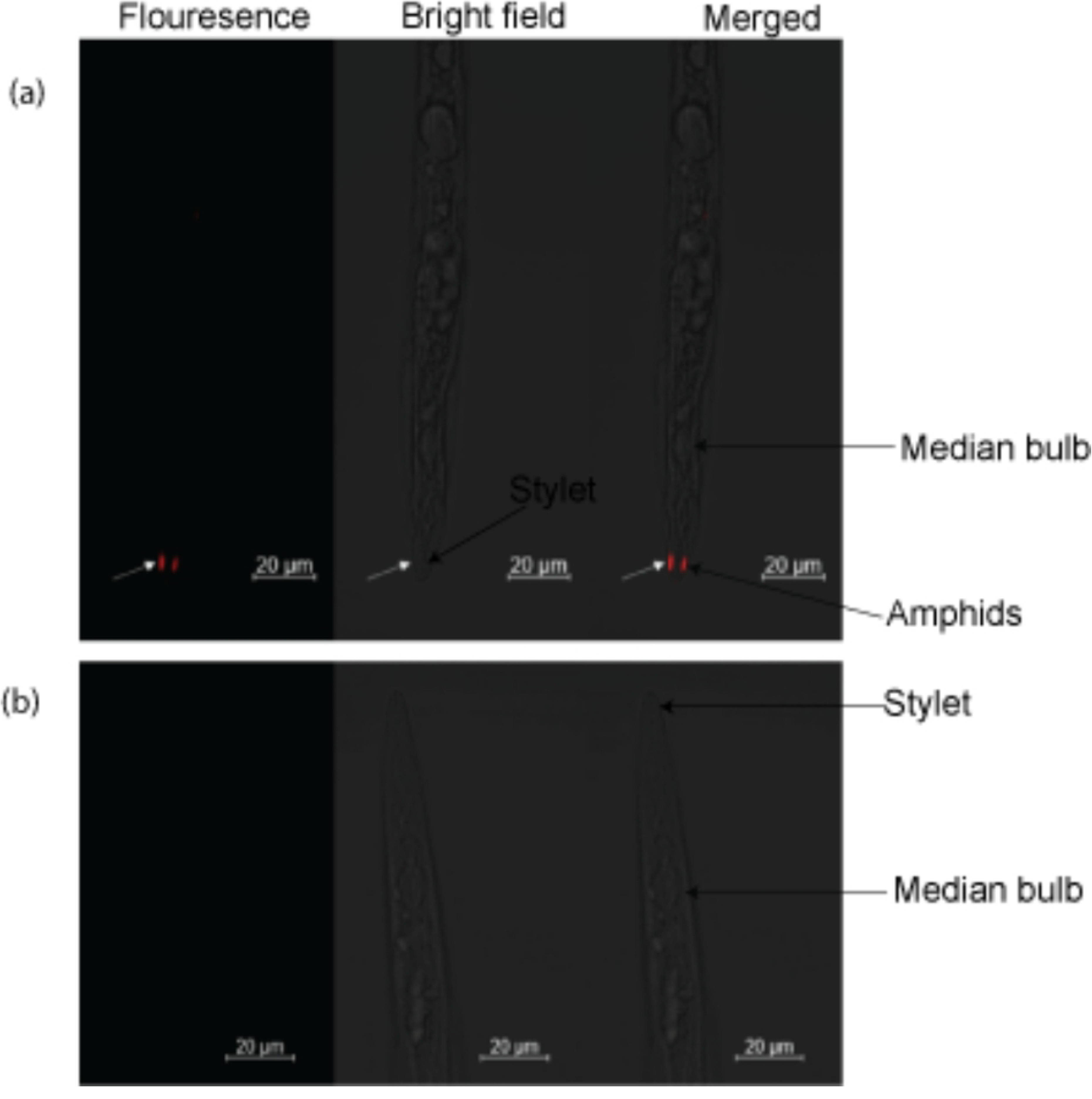
Fluorescence in situ hybridization of MjCRSP DNA probe conjugated to ATTO 647N dye. (a) DNA probes showing that transcripts of the gene were mainly detected (red staining) in the amphids secretory gland cells in the J2s of *M. javanica*. (b) No signal was detected for the control experiment.

## 3.4 Discussion

A long-standing question in plant pathology is how phytopathogens and pests, including PPNs, establish plant disease. Emerging evidence shows that PPNs have evolved mechanisms to evade, modulate, and suppress host immunity (Goverse & Mitchum, 2022, Siddique et al., 2022). The host-nematode interface, where molecular dialogues exist, determines the compatibility or incompatibility of the interaction (Siddique et al., 2022). In the quest for effective, long-lasting, and ecologically friendly management strategies, insights into the biology of these pathogenic microbes are essential. This entails identifying the molecular mechanisms developed by plant parasites to target host immunity and successfully colonize their host plants (Zhang et al., 2020).

In this instance, we undertook research on *M. javanica*, a species that shares evolutionary relationships with *M. incognita* (Castagnone-Sereno et al., 2013), an important agricultural pest that threatens world food security (Coyne et al., 2018). *M. javanica* is capable of infecting up to 770 plant species which include economically important food crops, vegetables, and weeds. (Mj distribution map and host range data: https://www.cabidigitallibrary.org/doi/10.1079/cabicompendium.33246).

Here, we identified a putative effector, MjCRSP, with a potential ICK structural motif that provides a high degree of thermal, chemical, and enzymatic stability provided by the disulfide bonds (Craik et al., 2001, Zhao et al., 2018, Park et al., 2014). ICK peptides have been linked to various multifunctional roles such as ion-blocking activity, insecticidal activity, and antimicrobial activity against bacteria and fungi (Park et al., 2014). Given the multiple functions of ICK peptides, there is a need for further studies to decipher additional functions associated with MjCRSP, outside of its involvement in the plant immune response, which was the focus of our study. Indeed, recent studies have demonstrated that depending on the developmental stage at which they are expressed, nematode effectors could serve numerous functions besides enhancing nematode virulence and countering plant defense. As an example, the BxICD1 effector from *Bursaphelenchus xylophilus* is involved in both nematode migration, which is essential for parasitism, and nematode pathogenicity (Li et al., 2024). Furthermore, Liu et al. (2023) reported that MiMIF-2 effector modifies the host rhizosphere microbiota to promote M. *incognita* parasitism.

A well-researched function of nematode effectors is their involvement in plant immunity. The identification of RKN effector proteins indispensable to nematode parasitism has been made possible largely by advances in omics technology and molecular research. MjTTL5 (Lin et al., 2016), Mj-NEROS (Stojilkovic et al., 2022), MjGO2 (Song et al., 2021b), and Mj-10A08 (Hu et al., 2022) are some of the well-known examples of *M. javanica* effectors that alter host immune responses. Since PCD inhibition is a potentially significant pathogenicity mechanism in biotrophic pests (Dalio et al., 2021), here we tested the cell-death-inducing and PCD inhibition properties of MjCRSP. Our infiltration assays show that MjCRSP does not induce PCD (**Figures 3 a1 and b2**) but consistently suppresses INF1-mediated cell death (**Figures 3 a2 and b2**). This indicates that MjCRSP evades plant immunity and can suppress INF1-mediated cell death immune signaling pathway. We also tested whether MjCRSP can suppress ETI-associated cell death following the co-expression of MjCRSP combined with ETI inducers (the cytoplasmic NBS-LRRs R3a and its cognate elicitor Avr3a) in *N. benthamiana* leaves. RKN effectors that have been shown to inhibit ETI-associated cell death include MiMsp40 (Niu et al., 2016) and Mj-10A08 (Hu et al., 2022). Our results show that MjCRSP failed to inhibit PCD induced by R3a/Avr3a cognate pairs (**Figure 3 a3 and b3**). It is important to note that pathogen effectors are selective on the immune-signaling pathways they manipulate. For instance, MiMsp40 was found to inhibit PCD triggered by R3a/Avr3a but not by Gpa2/RBP-1 immune signaling (Niu et al., 2016). Therefore, it would be interesting to establish if MjCRSP suppresses other ETI signalling pathways.

Besides manipulating PCD, RKN effectors have been reported to modulate immune responses such as ROS generation and accumulation of callose. For instance, MjTTL5 (Lin et al., 2016) and Mj-NEROS Stojilkovic et al. (2022) effectors interfere with ROS production while MIMSP40 suppresses callose deposition after elf18 PAMP treatment (Niu et al., 2016). In line with this, we evaluated the functionality of MjCRSP in manipulating ROS production and callose accumulation, which are components of the PTI (Zhao et al., 2020a). This was experimentally demonstrated by transiently expressing MjCRSP and a conserved Flg22 peptide derived from the bacterium flagellin. It is recognized by the *Arabidopsis* receptor kinase FLS2, resulting in the production of plant immunological responses like callose deposition, the expression of defense marker genes, ROS generation, and MAPK cascade activation (Zhao et al., 2020a). Compared with GFP/Flg22; MjCRSPp/Flg22 infiltrated leaves showed a significant reduction in callose deposition and ROS generation implying that MjCRSP can impede host PTI responses (**Figures 4a and b**). Both callose deposition and ROS are important downstream manifestations of PTI (Bentham et al., 2020, Lu & Tsuda, 2021). MjCRSP’s inhibition of ROS generation and callose levels may benefit nematode parasitism and reduce host disease resistance. Furthermore, low callose levels are essential for effective feeding and reproduction in nematodes (Xu *et al*., 2021). Thus, we deduced that MjCRSP is a secretory protein that functions as a novel effector of *M. javanica* and takes part in host-nematode interaction. Overall, plant defense mechanisms usually serve as a target for nematode effectors, which we infer also applies to the MjCRSP effector protein. MjCRSP undermines plant immune responses and based on the predicted NLS and transient expression analysis, it appears to function in the cytoplasm *of N. benthamiana* leaves (**Figures 5 and 6a**). However, the localisation of this effector in amphids is at odds with a role in the cytoplasm but rather, it would indicate a possible role in the host apoplast (Semblat *et al*., 2001, Eves-van den Akker *et al*., 2014). Thus, the mechanism of translocation of MjCRSP in the cytoplasm is subject to further investigation. Nonetheless, evidence shows that amphidial secretions might have diverse functions. A notable example of amphidial secretions functioning as a putative avirulence (Avr) factor in *M. incognita* is the protein MAP-1, whose release by pre-parasitic J2s indicated that it might be involved in the earliest phases of recognition between the nematodes and the plant (Semblat *et al*., 2001). This demonstrates even further the possibility that MjCRSP, based on our localization analysis, may serve a variety of activities that contribute to effective nematode parasitism and call for more research. A homolog of MjCRSP, Minc3s05519g38359 from *M. incognita* was discovered to be significantly expressed during the parasitic stages (Da Rocha et al., 2021) suggesting that this effector protein functions during RKN-host interactions.

## Conclusions

Based on the acquired data, we deduce that MjCRSP is a significant effector protein from *M. javanica* expressed in the amphids, and therefore in nematode secretory glands. Additionally, we showed that MjCRSP may promote nematode parasitism by inhibiting immune responses generated by PTI and PCD. Further research, such as host target investigation and pathologic cascade analysis, will elucidate the molecular mechanism underpinning MjCRSP in *M. javanica* parasitism.

## Supplementary information

Data generated or analysed during this study are available in the figshare repository, https://doi.org/10.6084/m9.figshare.30127405.v1.

## Notes

### Competing Interest Statement

The authors have declared no competing interest.

